# Dordaviprone/ONC201 Activation of the ClpP Mitochondrial Protease Inhibits the Growth of KRAS-Mutant Pancreatic Cancer and Overcomes RAS Inhibitor Resistance

**DOI:** 10.64898/2025.12.01.691471

**Authors:** Kristina Drizyte-Miller, Seamus E. Degan, Ryan D. Mouery, Amber M. Amparo, Brandon L. Mouery, Wen-Hsuan Chang, Runying Yang, Sheila R. Nicewarner Peña, Elisa Baldelli, Jeffrey A. Klomp, Edwin J. Iwanowicz, Lee M. Graves, Emanuel F. Petricoin, Adrienne D. Cox, Clint A. Stalnecker, Kirsten L. Bryant, Channing J. Der

## Abstract

Pancreatic ductal adenocarcinoma (PDAC) is characterized by KRAS-driven oncogenic signaling and tumor growth. Blockade of the KRAS ERK-MAPK pathway via small molecule direct RAS inhibitors has shown clinical promise, but intrinsic and acquired resistance limit the efficacy of these inhibitors as single agents. To identify potential combination strategies, we first assessed the ability of dordaviprone/ONC201, an FDA-approved agent, to inhibit PDAC cell and organoid growth. We observed that ONC201 reduced the growth of a broad panel of KRAS-mutant PDAC cell lines, and that the expression of mitochondrial protease ClpP was required for this efficacy. Mechanistically, we observed that treatment with ONC201 led to inhibition of mitochondrial respiration, causing a compensatory increase in glycolysis. Furthermore, ONC201 caused ClpP-dependent activation of PI3K-AKT-mTOR signaling and concurrent PI3K and mTOR inhibition further enhanced ONC201 growth suppression. ONC201 demonstrated an additive effect when combined with a RAS(ON) multi-selective inhibitor RMC-7977 in PDAC cells and organoids. Finally, PDAC cell lines with acquired resistance to RMC-7977 or *KEAP1* loss-driven resistance retained sensitivity to ONC201. We propose that concurrent treatment with ONC201 may delay onset of resistance to RAS inhibitor therapy.

**Statement of Significance:** ClpP activation by dordaviprone/ONC201 suppressed PDAC cell growth and overcame resistance to the RAS(ON) multi-selective inhibitor RMC-7977, providing support for investigating this combination as a potential combination treatment for KRAS-mutant pancreatic cancer.

## Introduction

Pancreatic ductal adenocarcinoma (PDAC) is among the deadliest forms of cancer, responsible for over 50,000 deaths annually in the United States (1). A defining feature of PDAC is the near-universal presence (>95%) of activating mutations in *KRAS*, typically at the G12, G13, or Q61 loci (2,3). One of the most profound consequences of oncogenic KRAS signaling is the rewiring of cellular metabolism, including altered oxidative phosphorylation and enhanced nutrient scavenging (4–6). These metabolic adaptations are not only a hallmark of PDAC but are also essential for tumor maintenance, making them a potential therapeutic vulnerability (7).

Direct pharmacologic inhibition of KRAS was long considered infeasible due to its structural properties (8). However, the identification of a previously undiscovered druggable pocket has led to a wave of new drugs targeting KRAS mutant-selective, pan-KRAS, and even pan-RAS isoforms (9–11). Several of these inhibitors have shown clinical promise, particularly as second-line therapies, with trials ongoing to explore their use in the first-line setting. Nevertheless, the emergence of resistance remains a major clinical challenge, underscoring the need for effective combination strategies (12).

Given the central role of KRAS in metabolic reprogramming, combining RAS-pathway inhibition with metabolic therapies is a rational approach. One such possibility is to leverage the mitochondrial dysfunction induced by dordaviprone/ONC201(13). Dordaviprone/ONC201 is a small-molecule imipridone that is FDA-approved for the treatment of H3K27-altered midline glioma (DMG) (14,15). Although ONC201 has shown remarkable efficacy across a broad range of tumor types, there is not clarity on the mechanisms underlying this efficacy. What is now clear is that ONC201 causes mitochondrial stress and disruption of glutamine metabolism (16,17).

Originally described as a dopamine receptor D2 (DRD2)-selective antagonist(18), new evidence has emerged that ONC201 is an equipotent activator of ATP-dependent Clp protease proteolytic subunit (ClpP), a mitochondrial matrix serine protease that regulates mitochondrial quality control through selective degradation of misfolded or damaged mitochondrial proteins (19). Additionally, ClpP degradation of respiratory complex proteins disrupts energy production (13,18,20). Following ONC201 treatment, ClpP proteolysis of respiratory complex components resulted in excessive production of reactive oxygen species (ROS) and subsequent cell death (21). Previous studies have also described ONC201 as an inducer of TRAIL-mediated apoptosis through antagonism of DRD2, but not all cell types demonstrate TRAIL-dependent anti-tumor effects (22). Induction of cell death by ONC201 as a single agent is also an open question, as anti-apoptotic activity has been evident only in combination with MEK inhibition (23). Currently, ONC201 is being evaluated as a dual ERK and AKT inhibitor in several settings including colorectal and endometrial cancer (24) (NCT05630794). However, another study shows ONC201 is a potent activator of PI3K-AKT signaling, and this signaling is required for ONC201 activity (15). Finally, ONC201 has shown efficacy in pre-clinical models of PDAC (25,26), but these studies do not comprehensively assess ONC201 and its potential combination with RAS inhibition.

In this study, we confirmed that ONC201 reduces PDAC cell growth in a ClpP-dependent manner. Through network-based analyses, we observed that ONC201 treatment enhanced RAS-PI3K-AKT signaling and induced mitochondrial dysfunction, contributing to additional metabolic stress. Based on these findings, we evaluated the combination of ONC201 with pan-RAS-targeted therapies and observed additive suppression of PDAC cell growth. We also generated KRAS-mutant PDAC cell lines that had acquired resistance to the RAS(ON) multi-selective inhibitor RMC-7977 and observed that ONC201 remained efficacious. Our results support the therapeutic potential of ClpP activation and suppression of mitochondrial function in combination with RAS inhibitors as an orthogonal combination treatment strategy for KRAS-mutant PDAC.

## Materials and Methods

### Cell culture

Human cell lines derived from patient-derived xenografts (PDX) (Pa01C, Pa02C, Pa14C, and Pa16C) were generously provided by Dr. Anirban Mitra (MD Anderson Cancer Center, Houston, TX). HEK 293T, MIA PaCa-2, PANC-1, and HPAC cell lines were obtained directly from ATCC. HPNE and HPNE-*KRAS^G12D^* cell lines were generated previously (27). All cell lines were grown in DMEM (Gibco, #11995065) supplemented with 10% fetal bovine serum (FBS) (Sigma) and 1% penicillin/streptomycin (Gibco). Cells were grown in a humidified chamber maintained at 37°C and 5% humidity. PDAC cell line identity was confirmed via short-tandem repeat (STR) sequencing in April 2017. Cells were periodically checked for *Mycoplasma* using the MycoAlert Kit (Lonza) and kept in culture for minimal passages.

Patient-derived KRAS-mutant PDAC organoids were provided by Dr. David Tuveson (Cold Spring Harbor Laboratory; hT105 and hM1A) and Dr. Calvin Kuo (Stanford University; PT3, PT6, and PT8), and were grown as previously described (28). Briefly, organoids were grown in growth factor–reduced Matrigel (Corning) domes in a custom media formulation containing: Advanced DMEM/F12 (Thermo Fisher Scientific)-based WRN conditioned medium [L-WRN (ATCC, CRL-3276)], 1× B27 supplement, 10 mmol/L HEPES, 0.01 μmol/L GlutaMAX (all from Thermo Fisher Scientific), 10 mmol/L nicotinamide (Sigma-Aldrich), 50 ng/mL hEGF (Peprotech), 100 ng/mL hFGF10 (Peprotech), 0.01 μmol/L hGastrin I (TOCRIS), 500 nmol/L A83-01 (TOCRIS), and 1.25 mmol/L (hM1A, hT105, hT106) or 1 mmol/L (PT3, PT6, PT8) N-acetylcysteine (Sigma-Aldrich). The medium for PT3, PT6, and PT8 also contained 10 μmol/L SB202190 (Sigma-Aldrich). Y27632 (Selleckchem, 10.5 μmol/L) was added for the first 2 days after reseeding single cells.

### Generation of RAS(ON) multi-selective inhibitor resistant cell lines

PDAC cell lines (∼20 x 10^6^) were plated in a 15 cm^2^ dish and treated with 100 nM RAS(ON) multi-selective inhibitor RMC-7977. Growth media were refreshed every 2-3 days with inhibitor. Cells were considered resistant after achieving at least five doublings in the presence of inhibitor.

### Chemical compounds

Dordaviprone (ONC201) (Selleckchem #S7963) and pictilisib (PI3Ki) (MedChemExpress #HY-50094) were obtained from commercial sources. TR107 was provided by Madera Therapeutics. The RAS(ON) multi-selective inhibitor RMC-7977 was provided by Revolution Medicines.

### Antibodies

Primary antibodies were purchased from Cell Signaling Technology: p44/42 MAPK (ERK1/2) (L34F12) (cat. #4696; RRID: AB_390780), phospho-p44/42 MAPK (pERK1/2) (Thr202/Tyr204) (D13.14.4E), Akt (pan) (40D4) (cat. # 2920), phospho-Akt (Ser473) (D9E) (cat. # 4060), GSK-3β (27C10) (cat. #9315), phospho-GSK-3β (Ser9) (cat. #9336), ACO2 (cat. #6922), CLPP (cat. #14181), GAPDH (D16H11) (cat. #5174), KEAP1 (cat. #8047), NQO1 (cat. #62262), and SLC7A11 (cat. #12691); Sigma-Aldrich: Vinculin (cat. #V9131; RRID:AB_477629) or from abcam: ALAS1 [EPR10247] (cat. # ab154860) and NRF2 (cat. #ab137550).

Secondary antibodies used were goat anti-rabbit IgG (H + L) cross-adsorbed secondary antibody, HRP (Invitrogen, #31462) and goat anti-mouse IgG (H + L) cross-adsorbed secondary antibody, HRP (Invitrogen, #31432).

Antibodies used for reverse-phase protein array (RPPA) analysis are listed in Supplementary Table 1.

### Immunoblotting

Cells were washed with ice cold PBS on ice and scraped using a standard Triton X-100 lysis buffer supplemented with phosphatase inhibitor cocktails I and II (Millipore; 524624 and 524625) plus a protease inhibitor cocktail (Roche; 11873580001). Lysates were kept on ice for 30 minutes with periodic vortexing before centrifugation at 12,700 rpm at 4°C for 10 minutes. The supernatant was then transferred to a new tube and protein levels were quantified using a Pierce BCA Protein Assay Kit (Thermo Fisher Scientific, #23225). For denaturation, Laemmli SDS sample buffer (4×; Bio-Rad, #1610747) supplemented with 0.1 mol/L dithiothreitol (Thermo Fisher Scientific, #R0862) was added to the lysates before incubation at 95°C for 5 minutes. Equal protein amounts were resolved by SDS-PAGE electrophoresis and transferred to polyvinylidene difluoride membranes (Millipore, IPVH00010) following standard immunoblotting procedures. The membranes were blocked in 3% BSA in TBS with 0.05% Tween 20 (TBST) for 1 hour. Membranes were then incubated with primary antibodies diluted in TBST at 4°C overnight. Thorough washing was performed prior to incubation with secondary antibodies diluted in TBST for 1 hour. Membranes were then washed again and developed by chemiluminescence (Bio-Rad ChemiDoc MP Imaging system).

### Cell proliferation

For 2D cell proliferation, cells were plated between 500 and 2,000 cells per well in a 96-well plate and allowed to attach and proliferate for 24 hours prior to treatment. The indicated inhibitors were added using the Tecan D300e Digital Dispenser. Following 5 days on treatment, cells were incubated with either calcein AM (Invitrogen, #C34852; final dilution 1:5,000) or Hoechst dye (Thermo Fisher Scientific, #H3570; final dilution 1:6,000) diluted in PBS. Plates were then incubated at room temperature for 10 to 20 minutes prior to readout. For calcein AM, fluorescence was measured on the MiniMax imager. For Hoechst staining, nuclei counting of Hoechst-stained cells was done via BioTek Cytation 1 Cell Imaging Multi-Mode Reader. Three biological and technical replicates were performed for each experiment. Cell counts were normalized to DMSO treated controls and percentage growth was calculated. GraphPad Prism (v10.3.1) was used to generate four-parameter dose–response curves and calculate AUC values.

For 3D organoid assays, 7,000 organoids were seeded per well of a 96-well plate with white walls and bottoms that were filled with 150 μL cold DMEM, 1:1,000 Y compound, 10% Matrigel. Following plating, organoids were allowed to settle for 48 hours before being treated for 6 days with the indicated inhibitor. Viability was measured using CellTiter-Glo 3D Cell Viability Assay (G9682) from Promega following manufacturer’s instruction. Representative images were acquired using the MiniMax imager.

Bliss synergy scores were determined using normalized cell counts and the SynergyFinder tool.

### Colony formation assay

Cells were plated at 1,000 -4,000 cells per plate and allowed to attach and proliferate 24 hours prior to treatment. Growth media supplemented with the inhibitor was added and allowed to grow until near maximal confluency was achieved in DMSO control plates. Inhibitor media was refreshed at 7 days. Once maximal confluence was achieved, plates were washed with PBS before being fixed with a crystal violet solution (0.05% crystal violet in 4% paraformaldehyde in 1× PBS) for 30 minutes. The solution was then aspirated and plates gently washed in water to remove excess before drying for 48 hours at room temperature. Images were taken on the Typhoon FLA 9500 Biomolecular Imager (GE Healthcare). Three biological replicates were performed unless otherwise indicated.

### *In vitro* CRISPR-Cas9 generation of *CLPP*-KO PANC-1 cells and sgKEAP1-KO cells

To generate PANC-1 cell lines with ClpP knockout, sgRNA sequences (2 control (*EGFP*) and 2 *CLPP)* were selected from the genome-scale CRISPR-Cas9 knockout screen (29). For cell lines with *KEAP1* knockdown, sgRNA sequences were selected from previously published *KEAP1* knockout studies (30). Sequences can be found in Supplementary Table 2. sgRNAs were amplified, and the amplified guides were ligated into the lentiCRISPRv2 puro backbone (Addgene, 52961) via NEBuilder HiFi DNA Assembly Master Mix (New England Biolabs, E2621). The ligation mix was then transformed into DH5α competent cells (50 μL, Thermo Fisher Scientific, EC0112). Clones were isolated and grown before DNA isolation using QIAGEN Plasmid *Plus* Midi Kit (Qiagen, 12143). HEK293T cells were grown overnight prior to DNA transfection using 3 μg psPAX2, 1 μg pMD2.G, 4 μg plasmid DNA in 400 μL Opti-MEM (Gibco), and 24 µL Fugene 6 (Promega, #E2691). The DNA mixture was incubated at room temperature for 15 minutes prior to dropwise addition to the HEK293T cells. Following an 18-hour incubation, the media was aspirated and replaced with DMEM supplemented with 20% FBS. Media with plasmid packaged in lentivirus was harvested 48 hours later and passed through a 0.45 µm filter attached to a syringe, aliquoted, and stored at -80°C. PANC-1 cells were plated in T25 flasks overnight, followed by transduction with 500 μL of virus containing plasmid DNA in 2 mL DMEM + 8 μg/mL polybrene. After 8 hours, transduction medium was replaced with normal DMEM. Following 48 hours, selection was initiated with puromycin.

### Oxygen consumption and extracellular acidification assay

To measure oxygen consumption rate (OCR) and extracellular acidification rate (ECAR), PDAC cell lines were plated into the XF96 cell culture microplate (Agilent #103794-100). Cells were then treated with 1 µM or 2.5 µM ONC201 for 24 hours. On the assay date, standard DMEM was replaced with assay medium containing 25 mM glucose, 1 mM glutamine, 1 mM sodium pyruvate and the respective drug treatment. OCR and ECAR were measured on an XF96 analyzer using the Seahorse Extracellular Flux Assay Kit (Agilent #103792-100). Results were normalized to cell number.

### Reverse phase protein array

PDAC cells were plated and treated with ONC201 for 24 and 72 hours. At the indicated time, plates were washed with cold PBS, snap frozen in liquid nitrogen, and stored at -80°C until all samples were collected. Each condition was performed in quadruplicate. RPPA was conducted as previously described (31). In brief, cell lysates were generated and spotted in triplicate (approximately 10 nL per spot) onto nitrocellulose-coated slides (Grace Biolabs, Bend, OR, USA) using a Quanterix 2470 Arrayer (Quanterix, Billerica, MA, USA). Control lysates were included on each slide to generate standard curves. The RPPA panel comprised 191 antibodies, each of which was validated for the presence of a single band at the expected molecular weight prior to use. Every slide was probed with one primary antibody. For secondary detection, either biotinylated goat anti-rabbit IgG (H+L) (1:7,500; Vector Laboratories, Burlingame, CA, USA) or rabbit anti-mouse IgG (1:10; DakoCytomation, Carpinteria, CA, USA) was applied. Signal amplification was performed using a tyramide-based avidin/biotin system (DakoCytomation), followed by labeling with streptavidin-conjugated IRDye 680 (LI-COR, Lincoln, NE, USA). Secondary-only incubations served as negative controls. Total protein was assessed using SpyroRed staining according to the manufacturer’s protocol (Molecular Probes, Eugene, OR, USA). Images were acquired with a Tecan PowerScanner (Tecan, Mannedorf, Switzerland) and quantified using MicroVigene software v5.1.0.0 (Vigenetech, Carlisle, MA, USA). For each sample, total protein levels were calculated by averaging the SpyroRed signal across replicate spots. Final intensities were obtained by subtracting negative-control signal from primary antibody signal, averaging the corrected values across triplicates, and normalizing to the sample’s total protein measurement.

Principal Component Analysis (PCA) was performed to evaluate consistency across replicates. All samples were retained for subsequent analysis. Antibodies with missing values or intensities under 1 were excluded from further analysis. Intensities were log_2_ transformed and median normalized. Differential expression was performed using the limma (V 3.21) package in R-studio (V 4.4.1). The “all” cell line condition was generated using each individual cell line as a blocking term. Plots were all generated using R-studio.

## Results

### ONC201 and TR107 activate CLPP to reduce PDAC cell growth

To evaluate the effectiveness of ONC201 to activate ClpP in *KRAS*-mutant PDAC, we treated a panel of KRAS-mutant PDAC cell lines with increasing doses of ONC201 for 24 hours. Consistent with increased ClpP protease activity, we observed a dose-dependent decrease in the abundance of mitochondrial proteins ALAS1 and ACO2, known targets of ClpP (Fig. 1A). We then expanded our panel of cell lines and tested the ability of ONC201 to suppress cell growth *in vitro*. ONC201 decreased 5-day cell proliferation in a dose-dependent manner across the entire panel (Fig. 1B).

**Figure 1.**
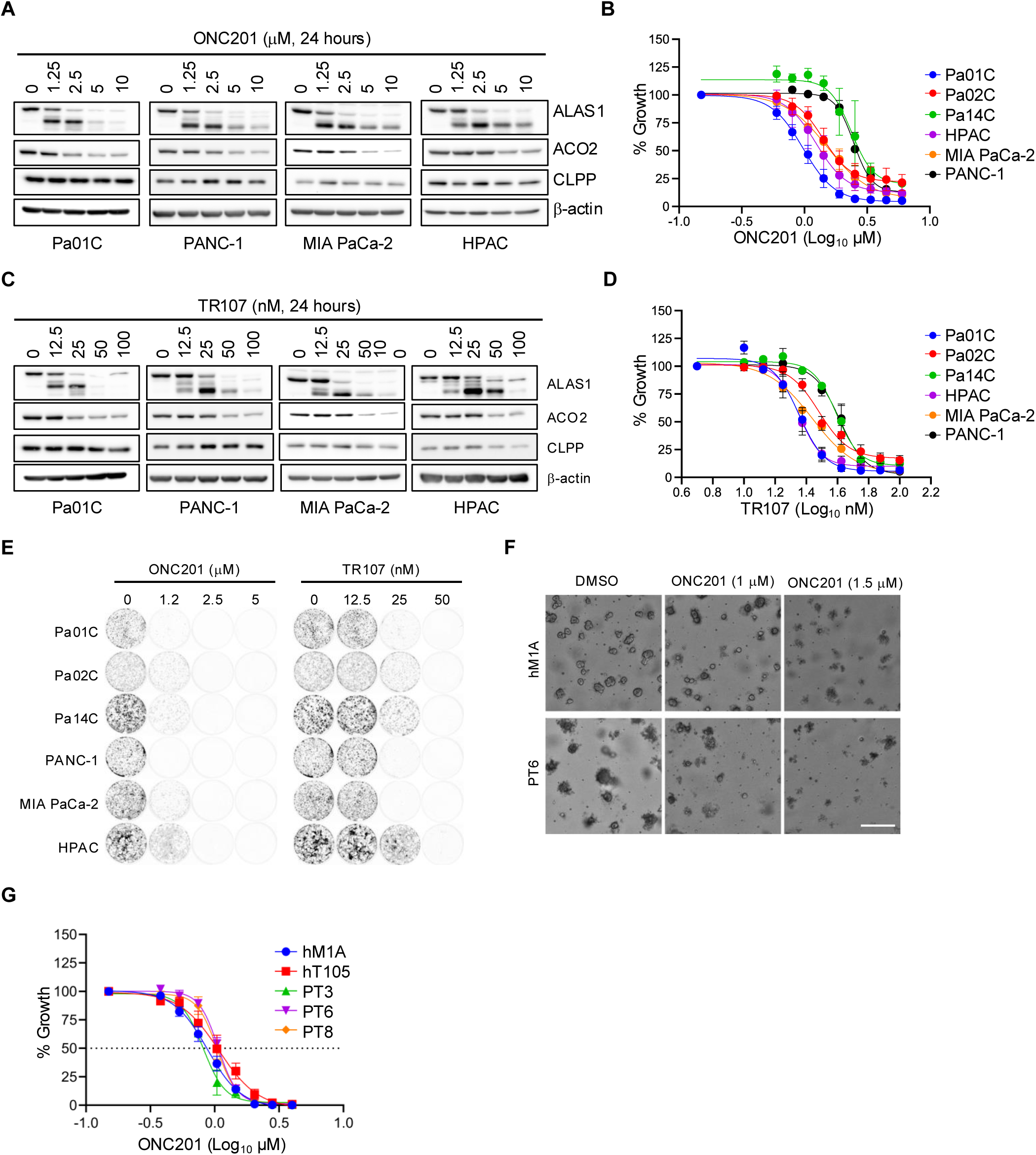
ONC201 and TR107 activate ClpP and reduce PDAC cell growth. **A, C,** Immunoblot for ALAS1, ACO2 and ClpP in *KRAS*-mutant PDAC cell lines treated for 24 hours with **A**, ONC201 or C, TR107. **B, D**, Proliferation following dose response of (**B**) ONC201 or (**D**) TR107 treatment for five days, represented as mean percent cell growth ± SEM (*n* = 3 independent experiments). **E,** 10- to 14-day clonogenic growth assay (images representative of *n* = 3). **F,** Representative images of hM1A and PT6 human PDAC patient-derived organoid (PDO cultures treated with DMSO or ONC201 (1 µM and 1.5 µM) for 6 days. Scale bars, 50 μm. **G,** Viability assay in PT3, PT6, PT8, hM1A and hT105 PDAC PDOs following increasing doses of ONC201 for 6 days.

To validate these findings, we performed the same assays using TR107, a small-molecule activator of ClpP that is chemically related to ONC201 (32). We observed similar activation of ClpP protease activity, as measured via immunoblotting (Fig. 1C) and decreased 5-day cell proliferation in the same panel of PDAC cell lines (Fig. 1D). To evaluate long-term growth suppression, we performed 10-14-day survival assays in the same panel of cell lines using both ONC201 and TR107. We observed robust reductions in colony formation across the panel of cell lines with both ONC201 and TR107 (Fig. 1E). To assess whether ClpP agonism is effective only in the context of mutant KRAS, we treated control or *KRAS*^G12D^-transformed human pancreatic ductal cells (HPNE). We observed no significant changes in 5-day proliferation between HPNE and HPNE-KRAS^G12D^ cells treated with either ONC-201 or TR107 (Supplementary Fig. S1A). These data show that an activating *KRAS*^G12D^ mutation has no effect on the efficacy of ONC201.

Finally, to test the ability of ONC201 to reduce growth in a more physiologically relevant setting, we used a panel of *KRAS*-mutant PDAC patient-derived organoids (PDOs) grown in 3D culture. Representative images of select PDOs illustrate their reduced viability following ONC201 treatment (Fig. 1F), with similar GI_50_ concentrations across the panel (Fig. 1G) as was also seen in 2D cell proliferation assays (Fig. 1D, Supplementary Fig. S1A). These findings show that ONC201 and a related ClpP agonist can reduce PDAC cell growth in both 2D and 3D conditions.

### ONC201 and TR107 reduce PDAC cell growth in a CLPP-dependent manner

Previous studies have identified ClpP and the dopamine receptor (DRD2) as direct targets of ONC201 (13,15,18). However, DRD2 is not detectable in *KRAS*-mutant PDAC, as demonstrated across multiple publicly available data sets (Fig. 2A). Although DRD3 may compensate for the lack of DRD2 in some situations (33,34), DRD3 is also not detectable in *KRAS*-mutant PDAC (Fig. 2A). To validate ClpP as the critical direct target of ONC201 in KRAS-mutant PDAC, we generated PANC-1 cell lines that stably express unique sgRNAs targeting ClpP. Genetic knockout of CLPP (Fig. 2B) resulted in loss of ClpP target inhibition upon treatment with ONC201 (Fig. 2B). Further, ONC201 was no longer able to inhibit proliferation in cells in the absence of ClpP (Fig. 2C). Using TR107 we observed a similar loss of target inhibition in ClpP knockout cells (Fig. 2D) and inability to reduce proliferation (Fig. 2E). Collectively, these findings identify ClpP as the critical target of ONC201 in *KRAS*-mutant PDAC and confirm its requirement for ONC201 efficacy.

**Figure 2.**
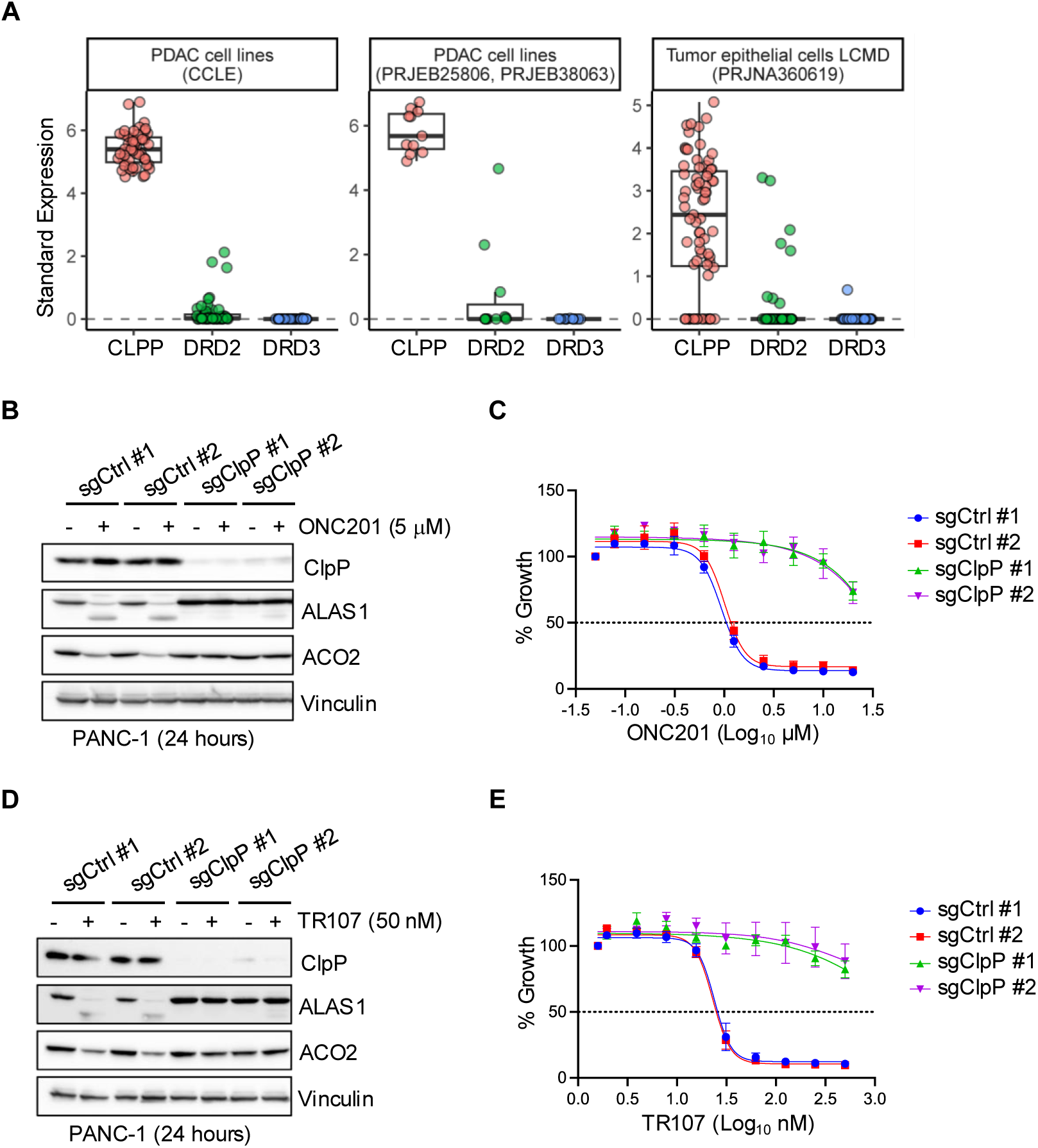
ONC201 and TR107 reduce PDAC cell growth in a ClpP-dependent manner. **A,** Graphs of expression levels of ClpP, DRD2, and DRD3 in publicly available PDAC cell line databases. **B, D**, Immunoblot for ClpP and its targets ALAS1 and ACO2 in ClpP-WT and ClpP-KO cell lines treated with (**B**) ONC201 or (**D**) TR107 for 24 hours. **C, E**, Proliferation following dose response of (**C**) ONC201 or (**E**) TR107 in ClpP-WT and ClpP-KO cell lines (five days), represented as mean percent cell growth ± SEM (*n* = 3 independent experiments).

### ONC201 inhibits mitochondrial respiration, resulting in a compensatory increase in glycolysis

Because of its role as a ClpP agonist, we hypothesized that treatment with ONC201 would lead to impaired mitochondrial respiration through oxidative phosphorylation (OXPHOS) and subsequently cause a switch to glycolytic metabolism. We first treated a panel of PDAC cell lines with ONC201 and used a Seahorse Assay to assess oxygen consumption rates (OCR), a measure of OXPHOS. We observed dose-dependent reductions in basal and maximal respiration, spare respiratory capacity, and ATP production (Fig. 3A), culminating in a complete collapse of mitochondrial respiration at the highest dose (Fig. 3B). The reduction in OCR was accompanied by decreased expression of all five OXPHOS complexes in a dose-dependent manner, as measured by immunoblotting with a human OXPHOS antibody cocktail (Fig 3C).

**Figure 3.**
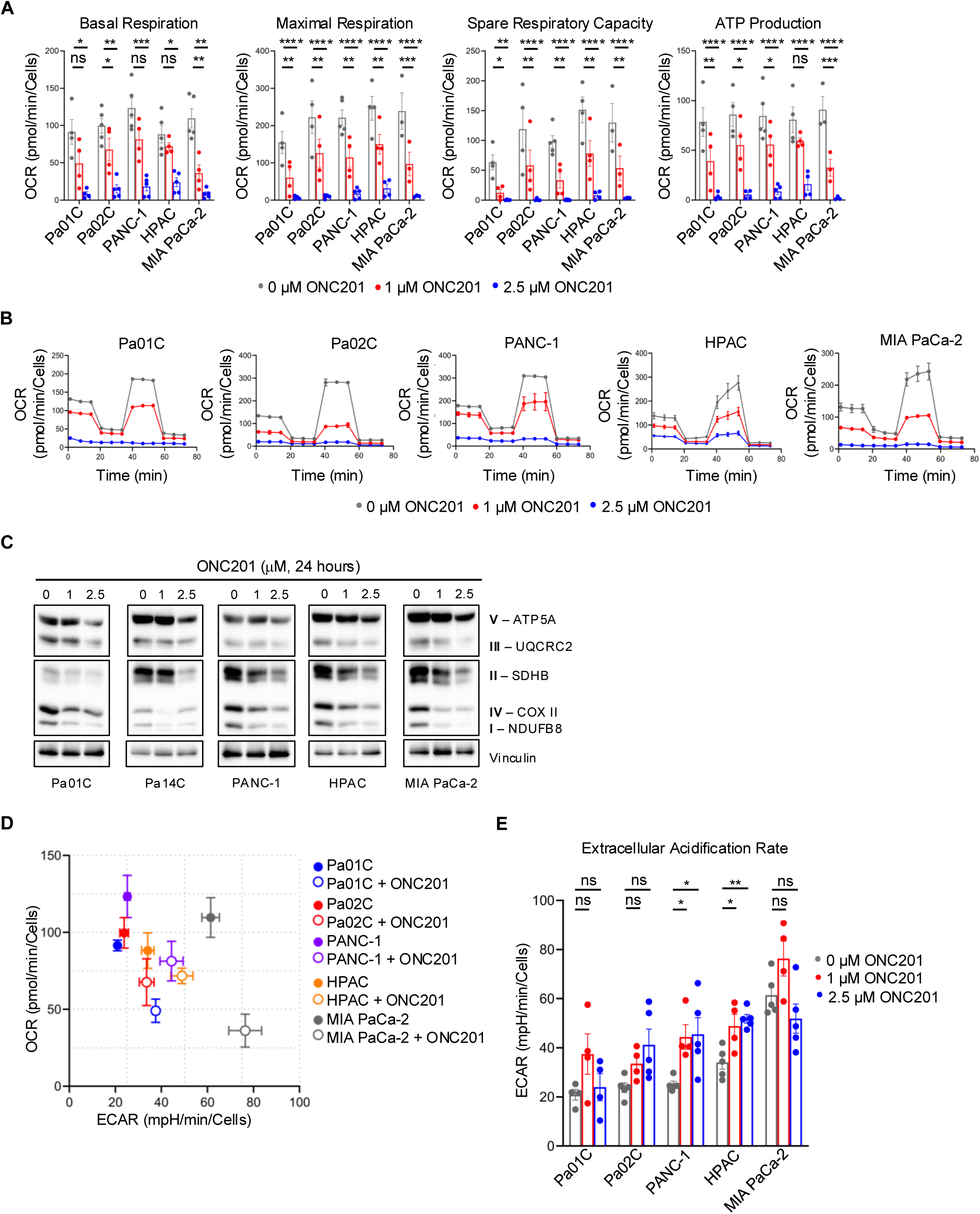
ONC201 inhibits mitochondrial respiration and causes a compensatory increase in glycolysis. **A,** Bar plots displaying OCR response of *KRAS*-mut PDAC cell lines treated with DMSO, 1 µM ONC201, or 2.5 µM ONC201 for 24 hours. P-values are from Dunnett’s multiple comparison test, comparing DMSO to treatment conditions. **B,** Rate plots displaying OCR over time detected by Seahorse Extracellular Flux Assay in cells treated as in **A**. **C,** Immunoblot for human OXPHOS antibody cocktail in cells treated as in **A**. **D,** Plot of OCR and ECAR measurements of *KRAS*-mutant PDAC cell lines treated with DMSO or 1 µM ONC201 (24 hours). Points represent geometric means ± SEM (*n* = 2) **E,** Bar plots displaying ECAR response of cells treated as in A. P-values are from Dunnett’s multiple comparison test, comparing DMSO to treatment conditions.

These results demonstrate that ONC201 blocks the ability to produce ATP through oxidative phosphorylation and points instead to a transition to an alternative process of ATP generation, i.e., glycolysis. To test whether treatment with ONC201 would lead to an increase in glycolysis in *KRAS*-mutant PDAC cells, we measured the extracellular acidification rate (ECAR), which increases as protons are generated during the conversion of glucose to lactate. We observed a shift across the panel of cell lines toward increased ECAR upon reduction in OCR (Fig. 3D). We validated that this increased ECAR occurs in a dose-dependent manner following ONC201 treatment (Fig 3E). These findings highlight that ONC201 disrupts PDAC cell metabolism, causing a collapse of OXPHOS and a compensatory increase in glycolysis.

### ONC201 treatment results in activation of the PI3K-AKT pathway

ONC201 is currently being evaluated as a dual PI3K and ERK pathway inhibitor in H3K27 mutant diffuse glioma/ diffuse intrinsic pontine glioma (DMG/DIPG) (24). However, whether this is true in the context of *KRAS*-mutant PDAC remains an open question. To investigate this, we performed reverse phase protein array (RPPA) analysis on our panel of PDAC cell lines following 24 and 72 hours of ONC201 treatment. We then performed differential antibody expression analysis and interrogated the changes. Globally, we observed few changes in the kinome across all cell lines and timepoints (Supplementary Fig. S4A). Although there were few signaling changes caused by ONC201 at 24 hours, levels of the PI3K downstream effectors pAKT S473 and pGSK3B S9 were significantly increased (Fig. 4A), and these increases were maintained at 72 hours post-treatment. We also observed significant increases in downstream effectors of AKT including pGSK3B, p4EBP1, and pBAD (Fig. 4A). When we isolated values from individual cell lines, we observed a general trend of increased PI3K-AKT signaling over time (Fig. 4B), as well as non-statistically significant increases in pERK signaling (Fig. 4B). Importantly, the increase in pAKT was ClpP-dependent, as ClpP-KO cells did not display increased pAKT, as indicated by immunoblotting (Supplementary Fig. S4B, C).

**Figure 4.**
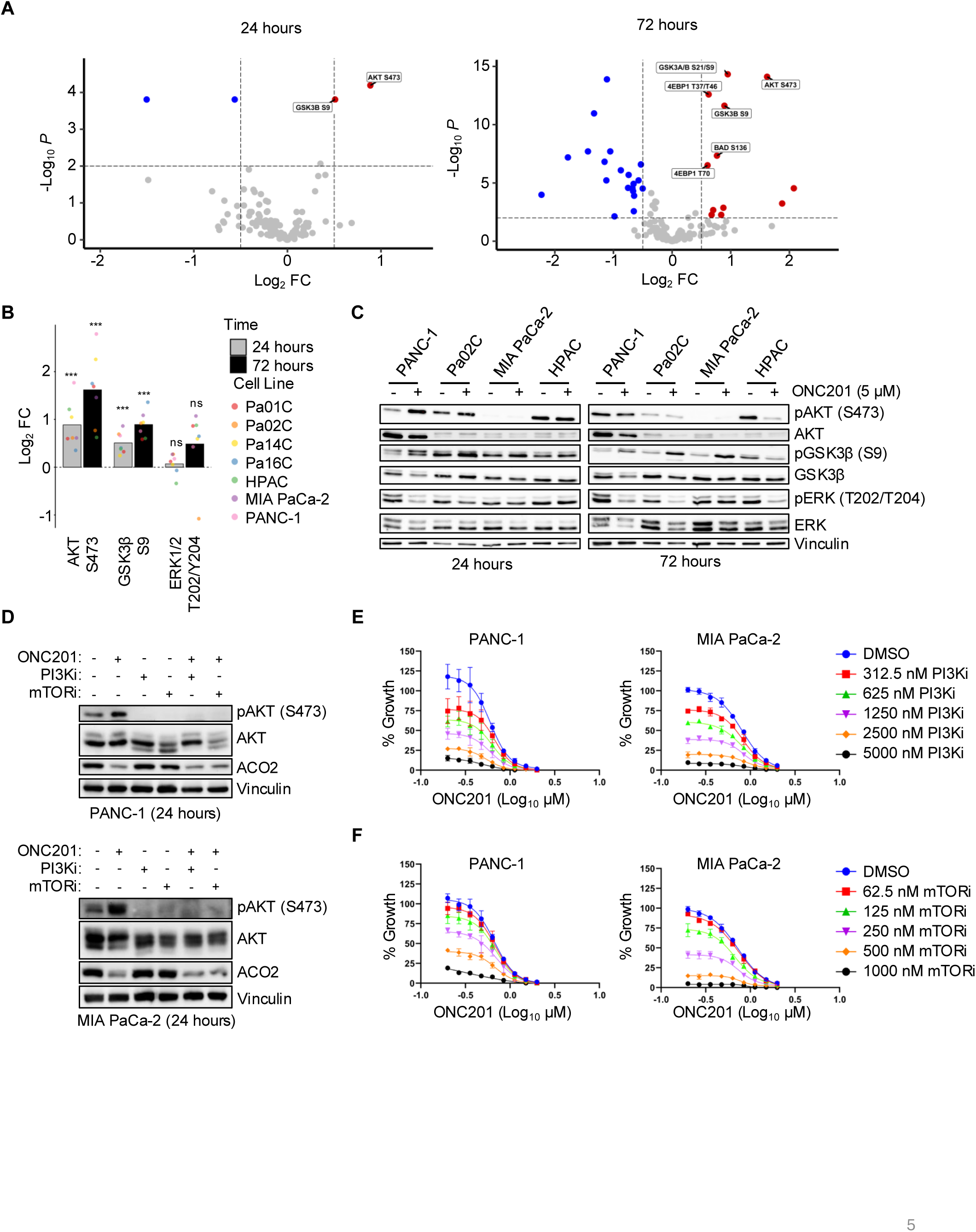
ONC201 treatment causes kinome reprogramming of the PI3K-AKT pathway, corresponding with CLPP activation. **A,** Volcano plots of differentially expressed proteins and phosphoproteins following 24 hours and 72 hours of ONC201 treatment. Significantly different proteins (*P* < 0.05, log_2_FC > 0.5) are annotated by blue (down-regulated) or red (up-regulated) points, with PI3K-AKT pathway components physically labeled. **B,** Bar plots of individual cell line log_2_FC values for selected proteins and phosphoproteins following 24 hours and 72 hours of ONC201 treatment. Statistical significance was determined by moderated *t*-tests from limma with Benjamini–Hochberg adjustment for multiple comparisons using the “all” cell line condition. **C,** Immunoblot for pAKT S473, AKT, pGSK3β S9, GSK3β, pERK 1/2 T202/Y204, and ERK 1/2 in select *KRAS*-mut PDAC cell lines treated with ONC201 for 24 hours and 72 hours. **D,** Immunoblot for pAKT S473, AKT, ClpP and its target ACO2 in MIA PaCa-2 and PANC-1 cells treated with (top) DMSO, ONC201, pictilisib (PI3Ki), or ONC201 plus pictilisib (24 hours) or (bottom) DMSO, ONC201, vistusertib (mTORi), or ONC201 plus vistusertib (24 hours). **E, F** Proliferation following five days of dose response treatment with (**E**) PI3Ki alone or with increasing doses of ONC201 or (**F**) mTORi alone or with increasing doses of ONC201, represented as mean percent cell growth ± SEM (*n* = 3 independent experiments).

We validated the RPPA findings by selecting a subset of cell lines and conducting immunoblotting under conditions that mirrored the RPPA assay. For example, we validated that pGSK3B and pAKT levels increased in response to ONC201 treatment at 24 hours, without significant changes in total GSK3B or total AKT (Fig 4C). Further, we observed minimal changes in pERK at 24 hours, again in line with our RPPA analysis. By 72 hours, we saw that levels of total AKT, ERK, and GSK3B decreased in ONC201-treated cells, whereas phosphorylated, active pAKT, pERK, and pGSK3B proteins increased. Our data challenges previous assertions about ONC201-induced signaling and points to a more nuanced reprogramming of the kinome following ONC201 treatment. Specifically, in *KRAS*-mutant PDAC cell lines, ONC201 is a PI3K-AKT pathway activator, not inhibitor.

Combination treatment of diffuse midline glioma with ONC201 and paxalisib, a pan-PI3K and mTOR inhibitor, was able to rescue increased PI3K signaling caused by ClpP-mediated mitochondrial degradation (15). To determine if PI3K or mTOR inhibition could reduce ONC201-induced pAKT activation in *KRAS*-mutant PDAC, we treated PANC-1 and MIA PaCa-2 cells with a PI3K or mTOR inhibitor alone or in combination with ONC201. Both PI3K and mTOR inhibitors reduced pAKT levels in ONC201-treated cells (Fig. 4D), which correlated with a reduction in 5-day proliferation (Fig. 4E). Collectively, we identify pAKT signaling as an on-target cellular stress response to ONC201 treatment in *KRAS*-mutant PDAC and demonstrate that treatment with inhibitors blocking this increase results in more potent inhibition of cellular proliferation.

### Concurrent treatment with ONC201 and the RAS(ON) multi-selective inhibitor RMC-7977 suppresses PDAC cell proliferation

The development of direct RAS inhibitors and their demonstrated efficacy herald an imminent shift in the treatment of *KRAS*-mutant PDAC from conventional cytotoxic agents to targeted therapies. Current and emerging targeted small molecules will be evaluated in combination with direct RAS inhibitors. Therefore, we assessed cellular proliferation following the combination of ONC201 and the RAS(ON) multi-inhibitor RMC-7977 (RASi), and found that the combination resulted in decreased proliferation as compared to single agent treatment (Fig. 5A). We then analyzed the combination’s effectiveness using SynergyFinder and determined that the effect of the combination was largely additive rather than synergistic (Fig. 5B). We extended the combination into organoid culture using our panel of *KRAS*-mutant PDAC organoids. Representative images of the organoids treated with single agent or the combination (Fig. 5C) illustrate that the reduced viability observed in 2D conditions with the combination of ONC201 and RASi also holds true in 3D conditions (Fig. 5D).

**Figure 5.**
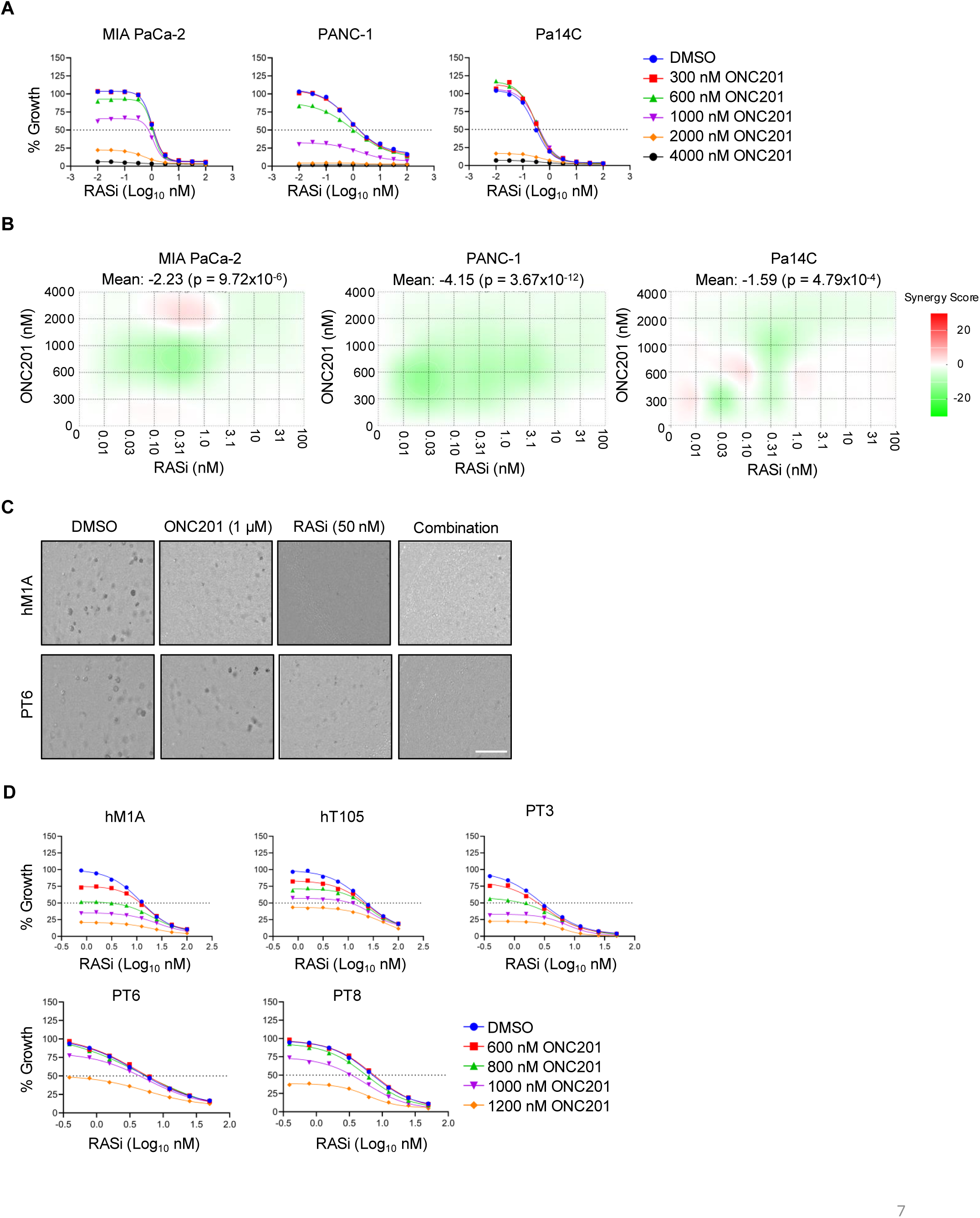
Combination of RAS(ON) multi-selective inhibitor and ONC201 reduces *KRAS*-mutant PDAC cell proliferation. **A,** Proliferation following five days of dose response treatment of *KRAS*-mutant PDAC cells with RMC-7977 (RASi) alone or with increasing doses of ONC201, represented as mean percent cell growth ± SEM (*n* = 3 independent experiments). **B,** Bliss synergy landscape plots for RASi and ONC201 in PANC-1, MIA PaCa-2, and Pa14C cells, with positive values (red) representing synergy, negative values (green) representing antagonism, and values near zero representing additive effects. **C,** Representative images of hM1A and PT6 human PDAC patient-derived organoid (PDO) cultures treated with DMSO or ONC201 (1 µM), RASi (50 nM) or the combination for 6 days. Scale bars, 50 μm. **D,** Viability assay in hM1A, hT105, PT3, PT6, and PT8 PDAC PDOs following increasing doses of RASi alone or with increasing doses of ONC201 for 6 days.

Despite the initial effectiveness of RAS inhibitor therapy, nearly all patients will develop resistance to RAS inhibitors through intrinsic or adaptive mechanisms (12). To understand the efficacy of ONC201 in the context of RASi-resistance (RASi-R), we generated pairs of RASi-R *KRAS*-mutant PDAC cell lines and their matched parental counterparts (Fig. 6A). We observed minimal changes in the GI_50_ values of ONC201 (Fig. 6B) and similar levels of target inhibition (Fig. 6C) in RASi-R cells when compared to parental cells, demonstrating that ONC201 is still efficacious in the setting of RASi resistance.

**Figure 6.**
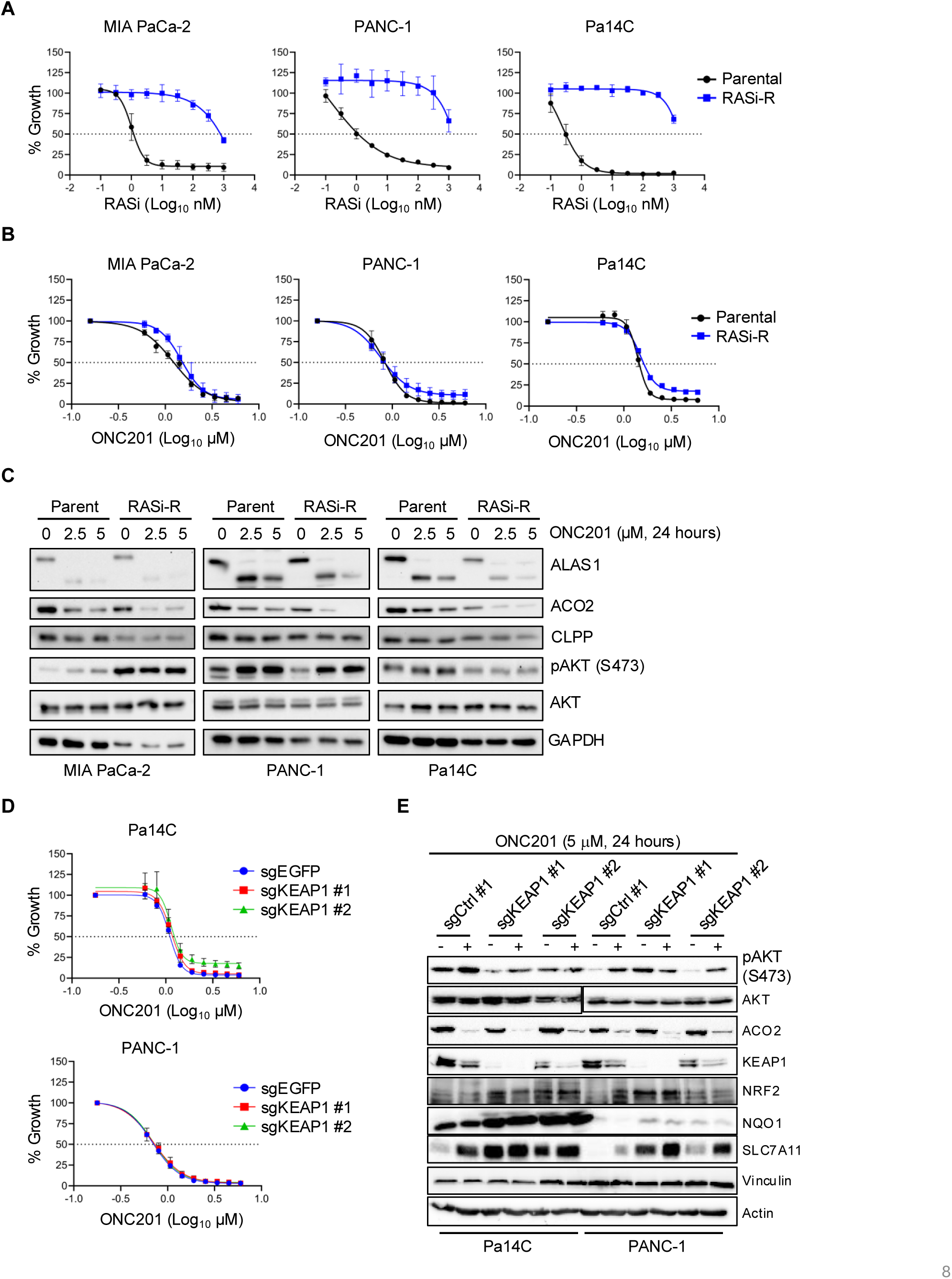
ONC201 reduces PDAC cell proliferation in the setting of RAS inhibitor resistance **A, B**, Proliferation following five days of dose response treatment with (**A**) RMC-7977 (RASi) or (**B**) ONC201 in parental cells or matched RAS inhibitor-resistant (RASi-R) cell lines, represented as mean percent cell growth ± SEM (*n* = 3 independent experiments). **C,** Immunoblot for ClpP and its targets ALAS1 and ACO2 and for pAKT S473 and AKT, in matched parental and RASi-R cell lines treated with increasing doses of ONC201 for 24 hours. **D**, Immunoblot of control (sgEGFP) and two independent (sgKEAP1#1, sgKEAP1#2) KEAP1 knockout cell lines for pAKT, ClpP target ACO2, KEAP1 and its targets NRF2, NQO1 and SLC7A11 following treatment with ONC201 for 24 hours. **E,** Proliferation of cells as in **D**, following five days of dose response treatment with ONC201, represented as mean percent cell growth ± SEM (*n* = 3 independent experiments).

Mutations in the Kelch-like ECH-associated protein 1 (*KEAP1*) gene have been shown to be a primary indicator of innate resistance to KRAS^G12C^ inhibitors in non-small cell lung cancer (NSCLC) (35). To understand the efficacy of RASi-resistance in a model of innate resistance, we generated PANC-1 and Pa14C cell lines that stably express unique sgRNAs targeting KEAP1. We saw minimal changes in GI_50_ values of ONC201 (Fig. 6D) and similar levels of target inhibition (Fig. 6E) in cells with genetic knockdown of KEAP1 compared to non-targeting controls. These findings support evaluation of ONC201 as a viable treatment strategy in the context of *KRAS*-mutant PDAC that become resistant to RAS(ON) inhibitors.

## Discussion

Effective targeted therapies for KRAS-mutant PDAC have been elusive (36). While mutant-selective inhibitors for *KRAS*^G12C^ have been approved, less than 1% of PDAC patients have this mutation (2). Even with growing evidence of the efficacy of mutant-selective *KRAS*^G12D^ and multi-RAS inhibitors, adaptive and intrinsic resistance remain major barriers to effective disease control (8,10). The recent FDA approval of dordaviprone/ONC201 in diffuse midline glioma demonstrates clinical benefit in cancer patients, but whether ONC201 is effective in *KRAS*-mutant PDAC in the context of RAS inhibition has been unknown (14,15). In this study, using a panel of PDAC cell lines and organoids, we demonstrate that ONC201 is not only broadly effective in inhibiting PDAC cell proliferation, but also shows additive growth inhibition when combined with the RAS(ON) multi-selective inhibitor RMC-7977. By using a panel of KRAS-mut PDAC cell lines and PDO’s, we demonstrate efficacy of ONC201 across a spectrum of secondary mutations beyond the activating KRAS mutation (Supplementary Table 3). This result suggests a broader use case for ONC201 across the spectrum of KRAS-mutant PDAC patients.

One of the largest open questions about ONC201 concerns the target(s) of the inhibitor. While it was initially described via machine learning-based algorithms as a DRD2 antagonist, further studies demonstrated that siRNA knockdown of DRD2 did not reduce ONC201’s growth inhibitory effect (18,37). In addition, future studies showed ClpP as a specific binding partner of ONC201 (13,16). Additionally, previous studies in PDAC cells and syngeneic mouse models did not assess whether DRD2 or ClpP were targets of ONC201 in this cancer type (25,26). In publicly available data, we found that pancreatic cancer cell lines do not express DRD2 or DRD3 at detectable levels, leaving only ClpP as a key target. Indeed, we found that ClpP protein was required for ONC201 to suppress PDAC cell growth, in line with previous studies demonstrating that ONC201 suppresses the growth of DIPG and triple negative breast cancer (TNBC) via ClpP agonism *in vitro* and *in vivo* (15,16).

ONC201 is currently under clinical investigation as a dual PI3K-AKT and ERK-MAPK pathway suppressor (38). However, although *KRAS*-mutant PDAC is largely dependent on ERK signaling (39), it was unclear whether the effects of ONC201 would be due to loss of ERK MAPK activity. Further, it was unknown how the lack of DRD2 will affect the signaling changes in response to ONC201. Studies have shown that DRD2 antagonism can drive the growth suppressive phenotype of ONC201 through RAS-ERK suppression in DMG cell lines and that ONC201 can synergize with the MEK inhibitor trametinib (20,23). Further, this DRD2 antagonism has been shown to be the primary mechanism of cell death induced by TRAIL signaling observed in multiple studies (24). In contrast to other suggestions (38), our findings support the hypothesis that ONC201 is a PI3K-AKT pathway activator rather than inhibitor, and that this activation is an on-target response to ClpP proteolysis. AKT phosphorylates GSK3β to inactivate it and promote cell survival (40). Further, GSK3β cooperates with KEAP1 to suppress NRF2 and the oxidative stress response (41). Oxidative stress has been reported to be induced in DMG following ONC201 treatment (15). In addition, we did not observe significant reduction in pERK levels at any point following ONC201 treatment. One potential hypothesis that emerges from these findings is that ClpP agonism is primarily responsible for PI3K-AKT pathway activation and that DRD2 antagonism is primarily responsible for RAS-ERK-MAPK pathway inactivation, as previously hypothesized (20).

The activating KRAS mutation has been shown to greatly alter PDAC cell metabolism, including mitochondrial dysfunction (42,43). This includes the enhanced uptake of glucose, glutamine, and the activation of nutrient scavenging pathways such as autophagy and macropinocytosis (44,45). Importantly, mitochondrial function and usage of oxidative phosphorylation is maintained within these tumors (46). We demonstrate that ONC201 can cause complete collapse of mitochondrial function and induce a further shift toward glycolysis. This supports previous findings in glioblastoma where treatment with ONC201 caused a similar loss of OXPHOS and increased glycolysis. (47). This finding points to a profound impact on *KRAS*-mutant PDAC cell metabolism in response to ONC201. Future studies may include assessment of glutamine dependence and nutrient scavenging pathways following ONC201 treatment.

Recent advances in the treatment of KRAS-mutant PDAC point to the field as on the precipice of a major shift toward the use of KRAS mutant-selective, pan-KRAS, and pan-RAS inhibitors (10,11,46). However, despite their success, intrinsic and acquired resistance remain major barriers to sustained disease control (12,47). Here we demonstrate that an orthogonal approach leveraging ONC201 and RMC-7977 can effectively suppress PDAC cell and PDO proliferation in both parental and RAS inhibitor-resistant models. With most acquired resistance mechanisms involving components of the RAS signaling network, a major focus of current efforts at establishing combinations have centered on more potently inhibiting the pathway through vertical RTK-RAS-ERK MAPK inhibition (48,49). ONC201 plus RAS inhibition represents a different type of combination, combining a metabolic therapy with RAS inhibition. Previous combinations with ONC201 centered around PI3K inhibition, with the hypothesis that suppressing the pro-survival PI3K-AKT signaling compensation caused by ONC201 would increase anti-tumor efficacy (14). Prior studies focused on using the blood-brain barrier-accessible PI3K inhibitor paxalisib in combination with ONC201 in DMG (15). However, paxalisib has been shown to be a dual PI3K and mTOR inhibitor (48). The demonstration that either PI3K or mTOR inhibition is sufficient to suppress ONC201-induced pAKT S473 levels and synergistically reduce PDAC cell proliferation strengthens the connection between ONC201 treatment and PI3K-AKT activation. The cooperation between the activating KRAS mutation and PI3K signaling in driving PDAC development suggests that adding ONC201 to proposed PI3K and RAS inhibitor combinations may prove to further enhance the anti-tumor efficacy of those combinations in KRAS-mutant PDAC. Overall, this study identifies a new potentially therapeutically tractable combination using ONC201 and RAS inhibition to more effectively target KRAS-mutant PDAC, even in the setting of RAS inhibitor resistance.

## Acknowledgments

The authors thank Dr. David Tuveson (Cold Spring Harbor Laboratory) and Dr. Calvin Kuo (Stanford University) for pancreatic organoid cultures. K. Drizyte-Miller was supported by National Cancer Institute (NCI) T32CA009156 and the American Cancer Society (ACS) PF-22-066-01-TBE. Support was provided by grants to A.D. Cox and/or C.J. Der from the NCI (R01CA42978, P50CA196510, P50CA257911, U01CA199235, P01CA203657 and R35CA232113), to C.J. Der from the Pancreatic Cancer Action Network (22-WG-DERB), and the Department of Defense (W81XWH2110692). K.L. Bryant was supported by Pancreatic Cancer Action Network/AACR grant 15-70-25-BRYA, NCI P50CA257911 and R37CA251877, and Department of Defense W81XWH2110693.

## Authors’ Contributions

**K. Drizyte-Miller:** Conceptualization, formal analysis, investigation, methodology, writing-review and editing. **S.E. Degan:** Formal analysis, investigation, methodology, writing-original draft, writing-review and editing. These authors contributed equally to the study. **R.D. Mourey:** formal analysis and investigation. **A.M. Amparo**: formal analysis and investigation. **R. Yang:** formal analysis, investigation. **S.R. Nicewarner Pena:** formal analysis, investigation. **E. Baldelli:** formal analysis, investigation. **L. M. Graves:** resources, supervision. **E.F. Petricoin:** resources, supervision, funding acquisition. **A.D. Cox:** Conceptualization, supervision, funding acquisition, writing-review and editing. **K.L. Bryant:** Conceptualization, formal analysis, resources, supervision, funding acquisition, writing-review and editing. **C.J. Der:** Conceptualization, formal analysis, resources, supervision, funding acquisition, writing-review and editing.

---

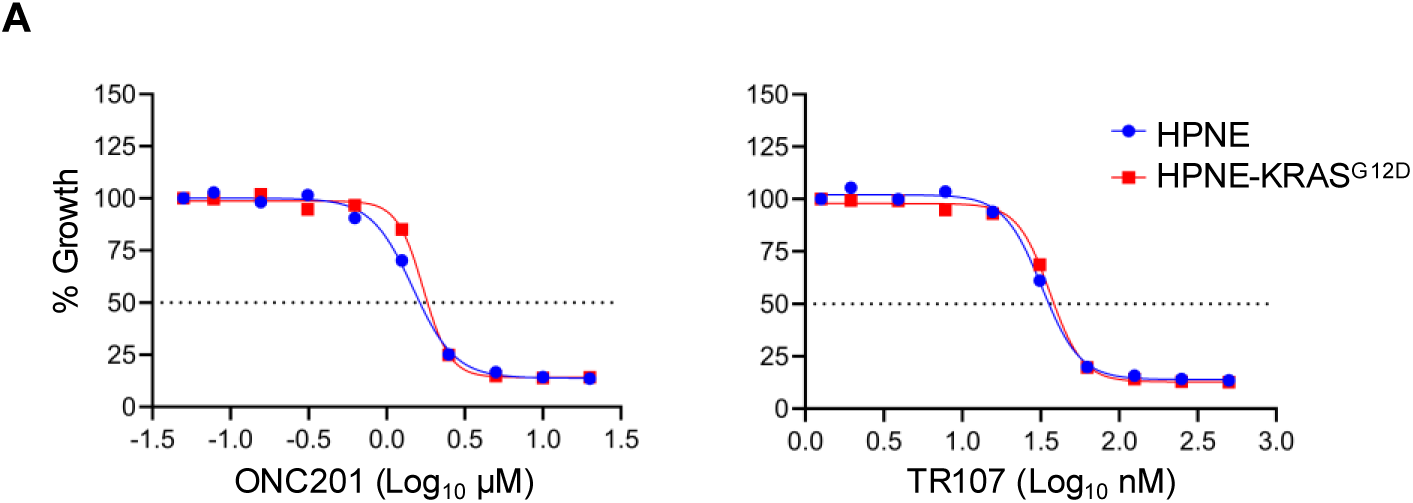

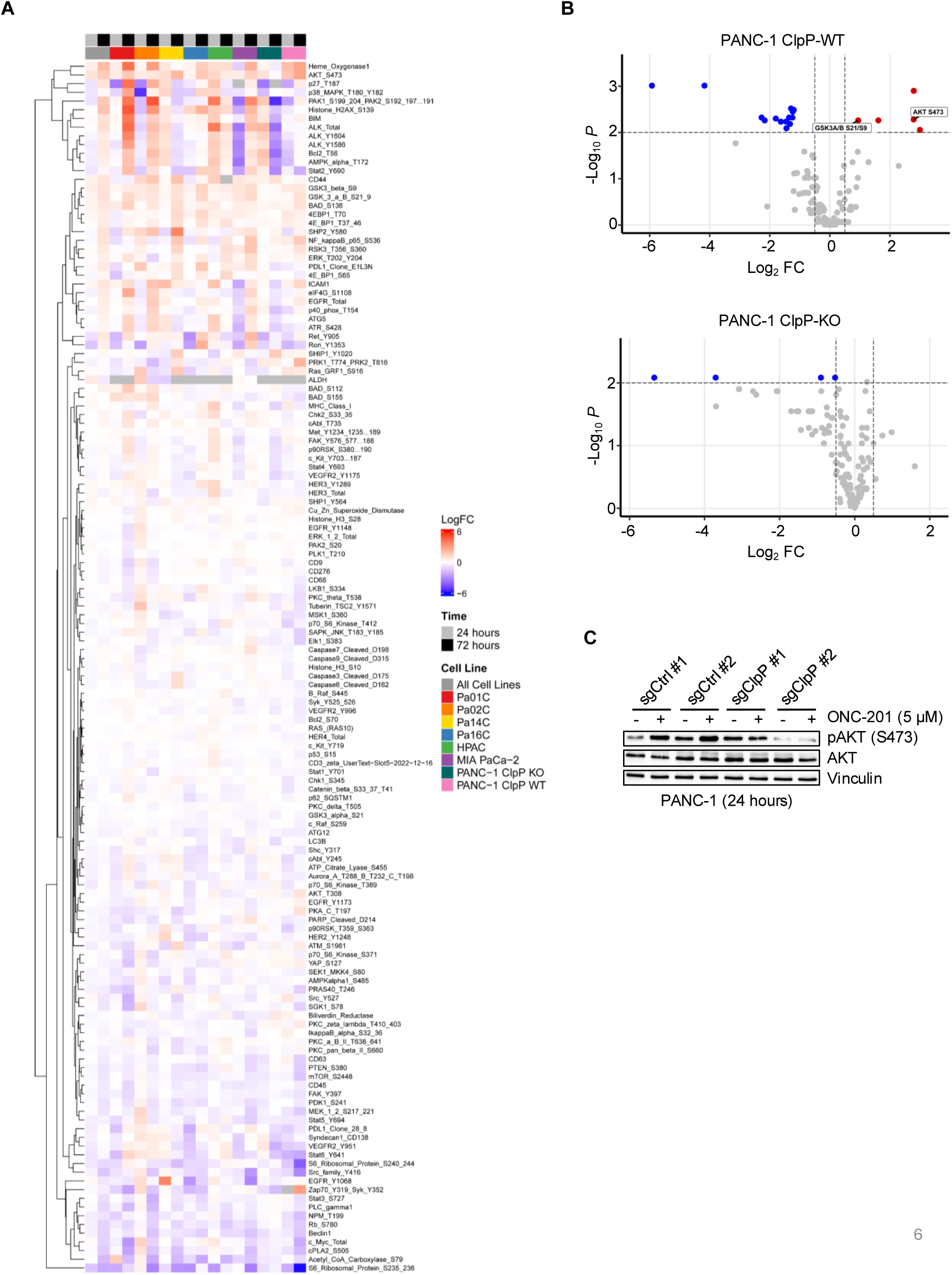

## REFERENCES

1. Siegel RL, Kratzer TB, Giaquinto AN, Sung H, Jemal A. Cancer statistics, 2025. CA Cancer J Clin. 2025;75:10–45.

2. Perelli L, Genovese G, Draetta GF. The KRAS mutational spectrum and its clinical implications in pancreatic cancer. Cancer Cell. Elsevier; 2024;42:1494–6.

3. McIntyre CA, Grimont A, Park J, Meng Y, Sisso WJ, Seier K, et al. Distinct clinical outcomes and biological features of specific KRAS mutants in human pancreatic cancer. Cancer Cell. Elsevier; 2024;42:1614–1629.e5.

4. Guo JY, Chen H-Y, Mathew R, Fan J, Strohecker AM, Karsli-Uzunbas G, et al. Activated Ras requires autophagy to maintain oxidative metabolism and tumorigenesis. Genes Dev. 2011;25:460–70.

5. Bryant KL, Stalnecker CA, Zeitouni D, Klomp JE, Peng S, Tikunov AP, et al. Combination of ERK and autophagy inhibition as a treatment approach for pancreatic cancer. Nat Med. 2019;25:628–40.

6. Yang A, Rajeshkumar NV, Wang X, Yabuuchi S, Alexander BM, Chu GC, et al. Autophagy Is Critical for Pancreatic Tumor Growth and Progression in Tumors with p53 Alterations. Cancer Discovery. 2014;4:905–13.

7. Kinsey CG, Camolotto SA, Boespflug AM, Guillen KP, Foth M, Truong A, et al. Protective autophagy elicited by RAF→MEK→ERK inhibition suggests a treatment strategy for RAS-driven cancers. Nat Med. Nature Publishing Group; 2019;25:620–7.

8. Cox AD, Fesik SW, Kimmelman AC, Luo J, Der CJ. Drugging the undruggable RAS: Mission Possible? Nat Rev Drug Discov. Nature Publishing Group; 2014;13:828–51.

9. Ostrem JM, Peters U, Sos ML, Wells JA, Shokat KM. K-Ras(G12C) inhibitors allosterically control GTP affinity and effector interactions. Nature. Nature Publishing Group; 2013;503:548–51.

10. Cregg J, Edwards AV, Chang S, Lee BJ, Knox JE, Tomlinson ACA, et al. Discovery of Daraxonrasib (RMC-6236), a Potent and Orally Bioavailable RAS(ON) Multi-selective, Noncovalent Tri-complex Inhibitor for the Treatment of Patients with Multiple RAS-Addicted Cancers. J Med Chem. American Chemical Society; 2025;68:6064–83.

11. Nokin M-J, Mira A, Patrucco E, Ricciuti B, Cousin S, Soubeyran I, et al. RAS-ON inhibition overcomes clinical resistance to KRAS G12C-OFF covalent blockade. Nat Commun. Nature Publishing Group; 2024;15:7554.

12. Ebright RY, Dilly J, Shaw AT, Aguirre AJ. Response and Resistance to RAS Inhibition in Cancer. Cancer Discov. 2025;15:1325–49.

13. Graves PR, Aponte-Collazo LJ, Fennell EMJ, Graves AC, Hale AE, Dicheva N, et al. Mitochondrial Protease ClpP is a Target for the Anticancer Compounds ONC201 and Related Analogues. ACS Chem Biol. American Chemical Society; 2019;14:1020–9.

14. Venneti S, Kawakibi AR, Ji S, Waszak SM, Sweha SR, Mota M, et al. Clinical Efficacy of ONC201 in H3K27M-Mutant Diffuse Midline Gliomas Is Driven by Disruption of Integrated Metabolic and Epigenetic Pathways. Cancer Discov. 2023;13:2370–93.

15. Jackson ER, Duchatel RJ, Staudt DE, Persson ML, Mannan A, Yadavilli S, et al. ONC201 in Combination with Paxalisib for the Treatment of H3K27-Altered Diffuse Midline Glioma. Cancer Res. 2023;83:2421–37.

16. Fennell EMJ, Aponte-Collazo LJ, Pathmasiri W, Rushing BR, Barker NK, Partridge MC, et al. Multi-omics analyses reveal ClpP activators disrupt essential mitochondrial pathways in triple-negative breast cancer. Front Pharmacol. 2023;14:1136317.

17. Fennell EMJ, Aponte-Collazo LJ, Graves PR, Herring LE, Iwanowicz EJ, Holmuhamedov E, et al. ONC201 and Its Potent Analogues Disrupt Mitochondrial Metabolic Function in Triple Negative Breast Cancer. The FASEB Journal. 2020;34:1–1.

18. Madhukar NS, Khade PK, Huang L, Gayvert K, Galletti G, Stogniew M, et al. A Bayesian machine learning approach for drug target identification using diverse data types. Nat Commun. 2019;10:5221.

19. Nouri K, Feng Y, Schimmer AD. Mitochondrial ClpP serine protease-biological function and emerging target for cancer therapy. Cell Death Dis. Nature Publishing Group; 2020;11:841.

20. Zhang Y, Tapinos N, Lulla R, El-Deiry WS. Dopamine pre-treatment impairs the anti-cancer effect of integrated stress response- and TRAIL pathway-inducing ONC201, ONC206 and ONC212 imipridones in pancreatic, colorectal cancer but not DMG cells. Am J Cancer Res. 2024;14:2453–64.

21. Ishizawa J, Zarabi SF, Davis RE, Halgas O, Nii T, Jitkova Y, et al. Mitochondrial ClpP-Mediated Proteolysis Induces Selective Cancer Cell Lethality. Cancer Cell. 2019;35:721–737.e9.

22. Ralff MD, Kline CLB, Küçükkase OC, Wagner J, Lim B, Dicker DT, et al. ONC201 Demonstrates Antitumor Effects in Both Triple-Negative and Non–Triple-Negative Breast Cancers through TRAIL-Dependent and TRAIL-Independent Mechanisms. Mol Cancer Ther. 2017;16:1290–8.

23. Lim B, Peterson CB, Davis A, Cho E, Pearson T, Liu H, et al. ONC201 and an MEK Inhibitor Trametinib Synergistically Inhibit the Growth of Triple-Negative Breast Cancer Cells. Biomedicines. Multidisciplinary Digital Publishing Institute; 2021;9:1410.

24. Allen JE, Krigsfeld G, Mayes PA, Patel L, Dicker DT, Patel AS, et al. Dual Inactivation of Akt and ERK by TIC10 Signals Foxo3a Nuclear Translocation, TRAIL Gene Induction, and Potent Antitumor Effects. Sci Transl Med. 2013;5:171ra17.

25. Jhaveri AV, Zhou L, Ralff MD, Lee YS, Navaraj A, Carneiro BA, et al. Combination of ONC201 and TLY012 induces selective, synergistic apoptosis in vitro and significantly delays PDAC xenograft growth in vivo. Cancer Biol Ther. 22:607–18.

26. Kumar V, Sethi B, Staller DW, Shrestha P, Mahato RI. Gemcitabine elaidate and ONC201 combination therapy for inhibiting pancreatic cancer in a KRAS mutated syngeneic mouse model. Cell Death Discov. Nature Publishing Group; 2024;10:158.

27. Campbell PM, Groehler AL, Lee KM, Ouellette MM, Khazak V, Der CJ. K-Ras Promotes Growth Transformation and Invasion of Immortalized Human Pancreatic Cells by Raf and Phosphatidylinositol 3-Kinase Signaling. Cancer Res. 2007;67:2098–106.

28. DeLiberty JM, Roach MK, Stalnecker CA, Robb R, Schechter EG, Pieper NL, et al. Concurrent Inhibition of the RAS-MAPK Pathway and PIKfyve Is a Therapeutic Strategy for Pancreatic Cancer. Cancer Res. 2025;85:1479–95.

29. Shalem O, Sanjana NE, Hartenian E, Shi X, Scott DA, Mikkelsen TS, et al. Genome-Scale CRISPR-Cas9 Knockout Screening in Human Cells. Science. American Association for the Advancement of Science; 2014;343:84–7.

30. Krall EB, Wang B, Munoz DM, Ilic N, Raghavan S, Niederst MJ, et al. KEAP1 loss modulates sensitivity to kinase targeted therapy in lung cancer. eLife. 6:e18970.

31. Bylicky MA, Shankavaram U, Aryankalayil MJ, Chopra S, Naz S, Sowers AL, et al. Multiomic-Based Molecular Landscape of FaDu Xenograft Tumors in Mice after a Combinatorial Treatment with Radiation and an HSP90 Inhibitor Identifies Adaptation-Induced Targets of Resistance and Therapeutic Intervention. Mol Cancer Ther. 2024;23:577–88.

32. Fennell EMJ, Aponte-Collazo LJ, Wynn JD, Drizyte-Miller K, Leung E, Greer YE, et al. Characterization of TR-107, a novel chemical activator of the human mitochondrial protease ClpP. Pharmacology Research & Perspectives. 2022;10:e00993.

33. Barroso-Chinea P, Luis-Ravelo D, Fumagallo-Reading F, Castro-Hernandez J, Salas-Hernandez J, Rodriguez-Nuñez J, et al. DRD3 (dopamine receptor D3) but not DRD2 activates autophagy through MTORC1 inhibition preserving protein synthesis. Autophagy. 16:1279–95.

34. Wang J-D, Cao Y-L, Li Q, Yang Y-P, Jin M, Chen D, et al. A pivotal role of FOS-mediated BECN1/Beclin 1 upregulation in dopamine D2 and D3 receptor agonist-induced autophagy activation. Autophagy. 2015;11:2057–73.

35. Tian L, Liu C, Zheng S, Shi H, Wei F, Jiang W, et al. KEAP1 mutations as key crucial prognostic biomarkers for resistance to KRAS-G12C inhibitors. J Transl Med. 2025;23:82.

36. Halbrook CJ, Lyssiotis CA, Pasca di Magliano M, Maitra A. Pancreatic cancer: Advances and challenges. Cell. 2023;186:1729–54.

37. Kline CLB, Ralff MD, Lulla AR, Wagner JM, Abbosh PH, Dicker DT, et al. Role of Dopamine Receptors in the Anticancer Activity of ONC201. Neoplasia. 2017;20:80–91.

38. Arrillaga-Romany I, Lassman A, McGovern SL, Mueller S, Nabors B, van den Bent M, et al. ACTION: a randomized phase 3 study of ONC201 (dordaviprone) in patients with newly diagnosed H3 K27M-mutant diffuse glioma. Neuro Oncol. 2024;26:S173–81.

39. Klomp JE, Diehl JN, Klomp JA, Edwards AC, Yang R, Morales AJ, et al. Determining the ERK-regulated phosphoproteome driving KRAS-mutant cancer. Science. 2024;384:eadk0850.

40. Doble B, Woodgett JR. GSK-3: tricks of the trade for a multi-tasking kinase. J Cell Sci. 2003;116:1175–86.

41. Baird L, Yamamoto M. The Molecular Mechanisms Regulating the KEAP1-NRF2 Pathway. Mol Cell Biol. 2020;40:e00099–20.

42. Bryant KL, Mancias JD, Kimmelman AC, Der CJ. KRAS: feeding pancreatic cancer proliferation. Trends Biochem Sci. 2014;39:91–100.

43. Kerk SA, Papagiannakopoulos T, Shah YM, Lyssiotis CA. Metabolic networks in mutant KRAS-driven tumours: tissue specificities and the microenvironment. Nat Rev Cancer. Nature Publishing Group; 2021;21:510–25.

44. Son J, Lyssiotis CA, Ying H, Wang X, Hua S, Ligorio M, et al. Glutamine supports pancreatic cancer growth through a KRAS-regulated metabolic pathway. Nature. 2013;496:101–5.

45. Weinberg F, Hamanaka R, Wheaton WW, Weinberg S, Joseph J, Lopez M, et al. Mitochondrial metabolism and ROS generation are essential for Kras-mediated tumorigenicity. Proc Natl Acad Sci U S A. 2010;107:8788–93.

46. Viale A, Pettazzoni P, Lyssiotis CA, Ying H, Sánchez N, Marchesini M, et al. Oncogene ablation-resistant pancreatic cancer cells depend on mitochondrial function. Nature. 2014;514:628–32.

47. Pruss M, Dwucet A, Tanriover M, Hlavac M, Kast RE, Debatin K-M, et al. Dual metabolic reprogramming by ONC201/TIC10 and 2-Deoxyglucose induces energy depletion and synergistic anti-cancer activity in glioblastoma. Br J Cancer. Nature Publishing Group; 2020;122:1146–57.

48. Heffron TP, Ndubaku CO, Salphati L, Alicke B, Cheong J, Drobnick J, et al. Discovery of Clinical Development Candidate GDC-0084, a Brain Penetrant Inhibitor of PI3K and mTOR. ACS Med Chem Lett. American Chemical Society; 2016;7:351–6.

