## Supplemental Files for "Dordaviprone/ONC201 Activation of the ClpP Mitochondrial Protease Inhibits the Growth of KRAS-Mutant Pancreatic Cancer and Overcomes RAS Inhibitor Resistance"

| Attribute | MIA PaCa-2 | Pa14C | Pa16C | PANC-1 | Pa01C | Pa02C | HPAC | PT3 | PT6 | PT8 | hT105 | hM1A |
|---|---|---|---|---|---|---|---|---|---|---|---|---|
| **KRAS** | G12C | G12D | G12D | G12D | G12D | Q61H | G12D | G12V | G12V | G12V | G12D | G12D |
| **p53** | R248W | Deleted | I255N | R273H | T155P | L247P | WT | R273H | G154V | Unknown | P142fs | R175H |
| **CDKN2A** | Deleted | Deleted | WT | Deleted | WT | Deleted | E120Ter | Unknown | Unknown | Unknown | Deleted | WT |
| **SMAD4** | WT | fs-indel | WT | WT | Deleted | WT | D52N_fs* | Unknown | R361H | Unknown | Deleted | WT |

| Antibody | Vendor | Catalogue Number | Dilution |
|---|---|---|---|
| 4E-BP1 (S65) | Cell Signaling Technologies | 9451 | 1:50 |
| 4E-BP1 (T37/46) | Cell Signaling Technologies | 9459 | 1:100 |
| Acetyl-CoA Carboxylase (S79) | Cell Signaling Technologies | 3661 | 1:50 |
| Akt (S473) | Cell Signaling Technologies | 9271 | 1:100 |
| Akt (T308) | Cell Signaling Technologies | 9275 | 1:100 |
| ALK | Cell Signaling Technologies | 3333 | 1:500 |
| ALK (Y1586) | Cell Signaling Technologies | 3348 | 1:500 |
| ALK (Y1604) | Cell Signaling Technologies | 3341 | 1:500 |
| AMPKalpha1 (S485) | Cell Signaling Technologies | 4184 | 1:1000 |
| AMPKalpha (T172) | Cell Signaling Technologies | 4188 | 1:1000 |
| Atg5 | Cell Signaling Technologies | 2630 | 1:1000 |
| ATP Citrate Lyase (S455) | Cell Signaling Technologies | 4331 | 1:1000 |
| ATR (S428) | Cell Signaling Technologies | 2853 | 1:500 |
| Aurora A (T288)/B (T232)/C (T198) | Cell Signaling Technologies | 2914 | 1:50 |
| BAD (S112) | Cell Signaling Technologies | 9291 | 1:200 |
| Bad (S136) | Cell Signaling Technologies | 9295 | 1:50 |
| Bad (S155) | Cell Signaling Technologies | 9297 | 1:100 |
| Bcl2 (T56) | Cell Signaling Technologies | 2875 | 1:1000 |
| Beclin 1 | Cell Signaling Technologies | 3738 | 1:100 |
| Biliverdin Reductase | Stressgen | OSA-400 | 1:500 |
| BIM | Cell Signaling Technologies | 2933 | 1:1000 |
| B-Raf (S445) | Cell Signaling Technologies | 2696 | 1:50 |
| c-Abl (T735) | Cell Signaling Technologies | 2864 | 1:50 |
| c-Abl (Y245) | Cell Signaling Technologies | 2861 | 1:100 |
| Caspase-3, cleaved (D175) | Cell Signaling Technologies | 9661 | 1:50 |
| Caspase-6, cleaved (D162) | Cell Signaling Technologies | 9761 | 1:50 |
| Caspase-7, cleaved (D198) | Cell Signaling Technologies | 9491 | 1:100 |
| Caspase-9, cleaved (D330) | Cell Signaling Technologies | 9501 | 1:50 |
| Catenin (beta) (S33/37/T41) | Cell Signaling Technologies | 9561 | 1:100 |
| CD68 | MyBiosource | MBS476527 | 1:500 |
| CD3ζ | BD | 556366 | 1:250 |
| CD44 | Cell Signaling Technologies | 3570 | 1:1000 |
| CD45 | BD | 610265 | 1:500 |
| CD63 | SantaCruz | Sc-5275 | 1:200 |
| CD276 | ProteinTech | 66481 | 1:3000 |
| CD9 | SantaCruz | Sc-13118 | 1:100 |
| Chk-1 (S345) | Cell Signaling Technologies | 2341 | 1:50 |
| Chk-2 (S33/35) | Cell Signaling Technologies | 2665 | 1:1000 |
| c-Kit (Y703) | Cell Signaling Technologies | 3073 | 1:1000 |
| c-Kit (Y719) | Cell Signaling Technologies | 3391 | 1:100 |
| c-Myc | Cell Signaling Technologies | 9402 | 1:100 |
| cPLA2 (S505) | Cell Signaling Technologies | 2831 | 1:1000 |
| Cu/Zn Superoxide Dismutase (SOD) | Stressgen | SOD-100 | 1:750 |
| 4EBP1 (T70) | Cell Signaling Technologies | 9455 | 1:1000 |
| EGFR (Y1068) | Cell Signaling Technologies | 2234 | 1:50 |
| EGFR (Y1148) | BioSource | 44-792 | 1:100 |
| EGFR (Y1173) | BioSource | 44-794 | 1:100 |
| EGFR | Cell Signaling Technologies | 2232 | 1:1000 |

| | | | |
|---|---|---|---|
| eIF4G (S1108) | Cell Signaling Technologies | 2441 | 1:1000 |
| Elk-1 (S383) | Cell Signaling Technologies | 9181 | 1:100 |
| ErbB2/HER2 (Y1248) | Imgenex | IMG-90189 | 1:500 |
| ErbB3/HER3 | Cell Signaling Technologies | 4754 | 1:1000 |
| ErbB3/HER3 (Y1289) (21D3) | Cell Signaling Technologies | 4791 | 1:200 |
| ErbB4/HER4 | Cell Signaling Technologies | 4795 | 1:1000 |
| ERK (T202/Y204) | Cell Signaling Technologies | 9101 | 1:1000 |
| ERK 1/2 | Cell Signaling Technologies | 9102 | 1:200 |
| FAK (Y397) | BD | 611806 | 1:1000 |
| FAK (Y576/577) | Cell Signaling Technologies | 3281 | 1:1000 |
| GSK-3α (S21) | Cell Signaling Technologies | 9337 | 1:2000 |
| GSK3β (S9) | Cell Signaling Technologies | 9336 | 1:1000 |
| GSK-3α/β (S21/9) | Cell Signaling Technologies | 9331 | 1:100 |
| Heme-Oxygenase-1 | Stressgen | SPA-894 | 1:500 |
| Histone H2AX (S139) | Cell Signaling Technologies | 9718 | 1:7500 |
| Histone H3 (S10) Mitosis Marker | Upstate | 06-570 | 1:200 |
| Histone H3 (S28) | Upstate | 07-145 | 1:1000 |
| IkappaBα (S32/36) | Cell Signaling Technologies | 9246 | 1:2000 |
| LC3B | Cell Signaling Technologies | 3868 | 1:1000 |
| LKB1 (S334) | Cell Signaling Technologies | 3055 | 1:50 |
| MEK1/2 (S217/221) | Cell Signaling Technologies | 9121 | 1:500 |
| Met (Y1234/1235) | Cell Signaling Technologies | 3126 | 1:200 |
| MHC Class I | SantaCruz | Sc-55582 | 1:100 |
| mTOR (S2448) | Cell Signaling Technologies | 2971 | 1:100 |
| MSK1 (S360) | Cell Signaling Technologies | 9594 | 1:1000 |
| NF-kappaB p65 (S536) | Cell Signaling Technologies | 3031 | 1:100 |
| NPM (T199) | Cell Signaling Technologies | 3541 | 1:1000 |
| p27 (T187) | Zymed | 71-7700 | 1:200 |
| p38 MAPK (T180/Y182) | Cell Signaling Technologies | 9211 | 1:100 |
| p40/phox (T154) | Cell Signaling Technologies | 4311 | 1:500 |
| p53 (S15) | Cell Signaling Technologies | 9284 | 1:1000 |
| p62/SQSTM1 (D5E2) | Cell Signaling Technologies | 8025 | 1:50 |
| p70 S6 Kinase (S371) | Cell Signaling Technologies | 9208 | 1:50 |
| p70 S6 Kinase (T389) | Cell Signaling Technologies | 9205 | 1:100 |
| p70 S6 Kinase (T412) | Upstate | 07-018 | 1:500 |
| p90RSK (T359/S363) | Cell Signaling Technologies | 9344 | 1:1000 |
| p90RSK (S380) | Cell Signaling Technologies | 9341 | 1:1000 |
| PAK1 (S199/204) PAK2 (S192/197) | Cell Signaling Technologies | 2605 | 1:1000 |
| PAK2 (S20) | Cell Signaling Technologies | 2607 | 1:1000 |
| PARP, cleaved (D214) | Cell Signaling Technologies | 9541 | 1:100 |
| PDK1 (S241) | Cell Signaling Technologies | 3061 | 1:1000 |
| PDL1 (Clone 28-8) | Abcam | Ab205921 | 1:500 |
| PDL1 (Clone E1L3N) | Cell Signaling Technologies | 13684 | 1:1000 |
| PKA C (T197) | Cell Signaling Technologies | 4781 | 1:1000 |
| PKC panβ II (S660) | Cell Signaling Technologies | 9371 | 1:1000 |
| PKCα/β II (T638/641) | Cell Signaling Technologies | 9375 | 1:1000 |
| PKCδ (T505) | Cell Signaling Technologies | 9374 | 1:1000 |
| PKCθ (T538) | Cell Signaling Technologies | 9377 | 1:1000 |
| PKCζ/ƛ (T410/403) | Cell Signaling Technologies | 9378 | 1:1000 |

| | | | |
|---|---|---|---|
| PLCɣ 1 | Cell Signaling Technologies | 2821 | 1:1000 |
| PLK1 (T210) | BD | 558400 | 1:1000 |
| PRAS40 (T246) | BioSource | 44-1100 | 1:1000 |
| PRK1 (T774)/2 (T816) | Cell Signaling Technologies | 2611 | 1:1000 |
| PTEN (S380) | Cell Signaling Technologies | 9551 | 1:500 |
| RAS (RAS10) | Cell Signaling Technologies | 05-516 | |
| c-Raf (S259) | Cell Signaling Technologies | 9421 | 1:100 |
| Rb (S780) | Cell Signaling Technologies | 3590 | 1:2000 |
| Ret (Y905) | Cell Signaling Technologies | 3221 | 1:100 |
| Ron (Y1353) | Epitomics | 5176-1 | 1:1000 |
| RSK3 (T356/S360) | Cell Signaling Technologies | 9348 | 1:500 |
| S6 Ribosomal Protein (S235/236) | Cell Signaling Technologies | 4856 | 1:200 |
| S6 Ribosomal Protein (S240/244) | Cell Signaling Technologies | 2215 | 1:1000 |
| SAPK/JNK (T183/Y185) | Cell Signaling Technologies | 9251 | 1:100 |
| SEK1/MKK4 (S80) | Cell Signaling Technologies | 9155 | 1:50 |
| SGK1 (S78) | Cell Signaling Technologies | 5599 | 1:500 |
| Shc (Y317) | Cell Signaling Technologies | 2431 | 1:500 |
| SHIP1 (Y1020) | Cell Signaling Technologies | 3941 | 1:1000 |
| SHP1 (Y564) | Cell Signaling Technologies | 8849 | 1:750 |
| SHP2 (Y580) | Cell Signaling Technologies | 5431 | 1:500 |
| Src family (Y416) | Cell Signaling Technologies | 2101 | 1:1000 |
| Src (Y527) | Cell Signaling Technologies | 2105 | 1:1000 |
| Stat1 (Y701) | Upstate | 07-307 | 1:1000 |
| Stat2 (Y690) | Cell Signaling Technologies | 4441 | 1:1000 |
| Stat3 (S727) | Cell Signaling Technologies | 9134 | 1:1000 |
| Stat4 (Y693) | Cell Signaling Technologies | 5267 | 1:1000 |
| Stat5 (Y694) | Cell Signaling Technologies | 9351 | 1:1000 |
| Stat6 (Y641) | Cell Signaling Technologies | 9361 | 1:1000 |
| Syk (Y525/526) | Cell Signaling Technologies | 2711 | 1:1000 |
| Syndecan1 (CD138) | Zymed | 36-2900 | 1:500 |
| Thioredoxin Reductase 1 (TrxR1) | SantaCruz | sc28321 | 1:50 |
| Tuberin TSC2 (Y1571) | Cell Signaling Technologies | 3614 | 1:1000 |
| Vav3 (Y173) | Biosource | 44-488 | 1:1000 |
| VEGFR2 (Y1175) | Cell Signaling Technologies | 2478 | 1:1000 |
| VEGFR2 (Y951) | Cell Signaling Technologies | 2471 | 1:1000 |
| VEGFR2 (Y996) | Cell Signaling Technologies | 2474 | 1:1000 |
| Vimentin | Cell Signaling Technologies | 3295 | 1:500 |
| YAP (S127) (D9W2I) | Cell Signaling Technologies | 13008 | 1:100 |
| Zap70 (Y319)/Syk (Y352) | Cell Signaling Technologies | 2701 | 1:500 |

| sgRNA | Sequence | Source |
|---|---|---|
| sgEGFP #1 | GGGCGAGGAGCTGTTCACCG | PMCID: PMC4089965 |
| sgEGFP#2 | GAGCTGGACGGCGACGTAAA | PMCID: PMC4089965 |
| sgCLPP #1 | GCAGCGGTGCCTGCACGCGA | PMCID: PMC4089965 |
| sgCLPP #2 | GCGCCTATGACATCTACTCG | PMCID: PMC9354705 |
| sgKEAP1 #1 | CTTGTGGGCCATGAACTGGG | PMCID: PMC5305212 |
| sgKEAP1 #2 | GTGTCCTCCACGTCATGAA | PMCID: PMC5305212 |
